# SARS-CoV-2 variant transition dynamics are associated with vaccination rates, number of co-circulating variants, and natural immunity

**DOI:** 10.1101/2022.11.18.517139

**Authors:** Lauren J Beesley, Kelly R Moran, Kshitij Wagh, Lauren A Castro, James Theiler, Hyejin Yoon, Will Fischer, Nick W Hengartner, Bette Korber, Sara Y Del Valle

**Affiliations:** Statistical Sciences, Los Alamos National Laboratory, Los Alamos, NM, USA; Theoretical Biology and Biophysics, Los Alamos National Laboratory, Los Alamos, NM, USA; Space Data Science and Systems, Los Alamos National Laboratory, Los Alamos, NM, USA; Center for Nonlinear Studies, Los Alamos National Laboratory, Los Alamos, NM, USA; The New Mexico Consortium, Los Alamos, New Mexico, USA; Information Systems and Modeling, Los Alamos National Laboratory

**Keywords:** SARS-CoV-2, COVID-19, variant transition, GISAID

## Abstract

**Background:** Throughout the COVID-19 pandemic, the SARS-CoV-2 virus has continued to evolve, with new variants outcompeting existing variants and often leading to different dynamics of disease spread.

**Methods:** In this paper, we performed a retrospective analysis using longitudinal sequencing data to characterize differences in the speed, calendar timing, and magnitude of 13 SARS-CoV-2 variant waves/transitions for 215 countries and sub-country regions, between October 2020 and October 2022. We then clustered geographic locations in terms of their variant behavior across all Omicron variants, allowing us to identify groups of locations exhibiting similar variant transitions. Finally, we explored relationships between heterogeneity in these variant waves and time-varying factors, including vaccination status of the population, governmental policy, and the number of variants in simultaneous competition.

**Findings:** This work demonstrates associations between the behavior of an emerging variant and the number of co-circulating variants as well as the demographic context of the population. We also observed an association between high vaccination rates and variant transition dynamics prior to the Mu and Delta variant transitions.

**Interpretation:** These results suggest the behavior of an emergent variant may be sensitive to the immunologic and demographic context of its location. Additionally, this work represents the most comprehensive characterization of variant transitions globally to date.

**Funding:** Laboratory Directed Research and Development (LDRD), Los Alamos National Laboratory

**Research in context:** *Evidence before this study:* SARS-CoV-2 variants with a selective advantage are continuing to emerge, resulting in variant transitions that can give rise to new waves in global COVID-19 cases and changing dynamics of disease spread. While variant transitions have been well studied individually, more work is needed to better understand how variant transitions have occurred in the past and how properties of these transitions may relate to vaccination rates, natural immunity, and population demographics.

*Added value of this study:* Our retrospective study integrates metadata based on 12.8 million SARS-CoV-2 sequences available through the Global Initiative on Sharing All Influenza Data (GISAID) with clinical and demographic data to characterize heterogeneity in variant waves/transitions across the globe throughout the COVID-19 pandemic. We demonstrate that properties of the variant transitions (e.g., speed, timing, and magnitude of the transition) are associated with vaccination rates, prior COVID-19 cases, and the number of co-circulating variants in competition.

*Implications of all the available evidence:* Our results indicate that there is substantial heterogeneity in how an emerging variant may compete with other viral variants across locations, and suggest that each location’s contemporaneous immunologic landscape may play a role in these interactions.

## 1 Introduction

Since the first SARS-CoV-2 viral sequence became available in January of 2020 ^1^, there have been over 630 million confirmed cases of COVID-19 globally^2^, leading to over 6.5 million deaths. SARS-CoV-2 is continuously evolving, and global transitions to newly emergent variants can generate waves of disease spread. The selective advantage of a new variant over existing variants is often associated with increased infectivity (e.g., through enhanced receptor binding or spike processing) and/or increased resistance to neutralizing antibodies induced by vaccination, prior infection, or both.^3,4,5,6,7,8,9,10^ Prior infections with different variants can be associated with differing protection against newly emergent variants,^11^ and we hypothesize that *vaccination rates and the history of prior infecting variants may impact the rate at which an emerging variant out-competes existing variants* to become the dominant form of the virus in a given country or state.

To explore this hypothesis and characterize heterogeneity in the speed, timing, and magnitude of variant transitions globally, we performed a retrospective analysis of over 12.8 million SARS-CoV-2 sequences reported to the Global Initiative on Sharing All Influenza Data (GISAID) between October 2020 and October 2022.^12^ We used multinomial regression spline modeling to estimate and summarize variant transition dynamics across 215 countries and sub-country regions and 13 SARS-CoV-2 variant waves. In a sub-analysis, we also characterize recently emergent Omicron sub-lineages BA.2.75, XBB.1/XBB, and BQ.1^10^ using GISAID data collected through November 3rd, 2022. Our results illustrate large heterogeneity in variant transitions between locations. For Omicron, we clustered geographic locations in terms of their variant behavior, allowing us to identify groups of locations with similar transition dynamics. We then leveraged clinical and demographic data to explore how properties of variant waves relate to time-varying factors, including vaccination status of the population, governmental policy, and the number of variants in competition. This work demonstrates an association between the behavior of an emerging variant and the immunologic and demographic context of the population. Additionally, this work represents the most comprehensive characterization of SARS-CoV-2 variant transitions globally to date.

## 2 Methods

### 2.1 Data sourcing and processing

Analyzed data streams (summarized in **Supp. Figure A.1**) are described below. All data were aggregated by date and location. We defined spatial locations at the country level and, for select countries having sufficient data, the sub-country region level.

#### GISAID SARS-CoV-2 data

Data for over 12.8 million SARS-CoV-2 sequences reported to GISAID by 10/01/2022 (https://gisaid.org) were obtained through the COVID-19 Viral Genome Analysis Pipeline https://cov.lanl.gov. We resolved location name inconsistencies and removed sequences with evident entry errors. We then categorized the sequences by variant (e.g., Alpha, Delta) based on each sequence’s Pango nomenclature SARS-CoV-2 lineage designation^13^, after excluding records designated “None” or “Unassigned”. Pango sub-lineage groupings as of 10/01/2022 are provided in **Supp. Table A.2**. **Figure 1** illustrates the reported variant proportions over time globally and for four example countries. In an emerging variants sub-analysis focusing on Omicron BA.2.75, XBB, and BQ.1 Pango lineages (defined in **Supp. Table A.2**), we used updated GISAID data with sequences reported by 11/03/2022.

**Figure 1:**
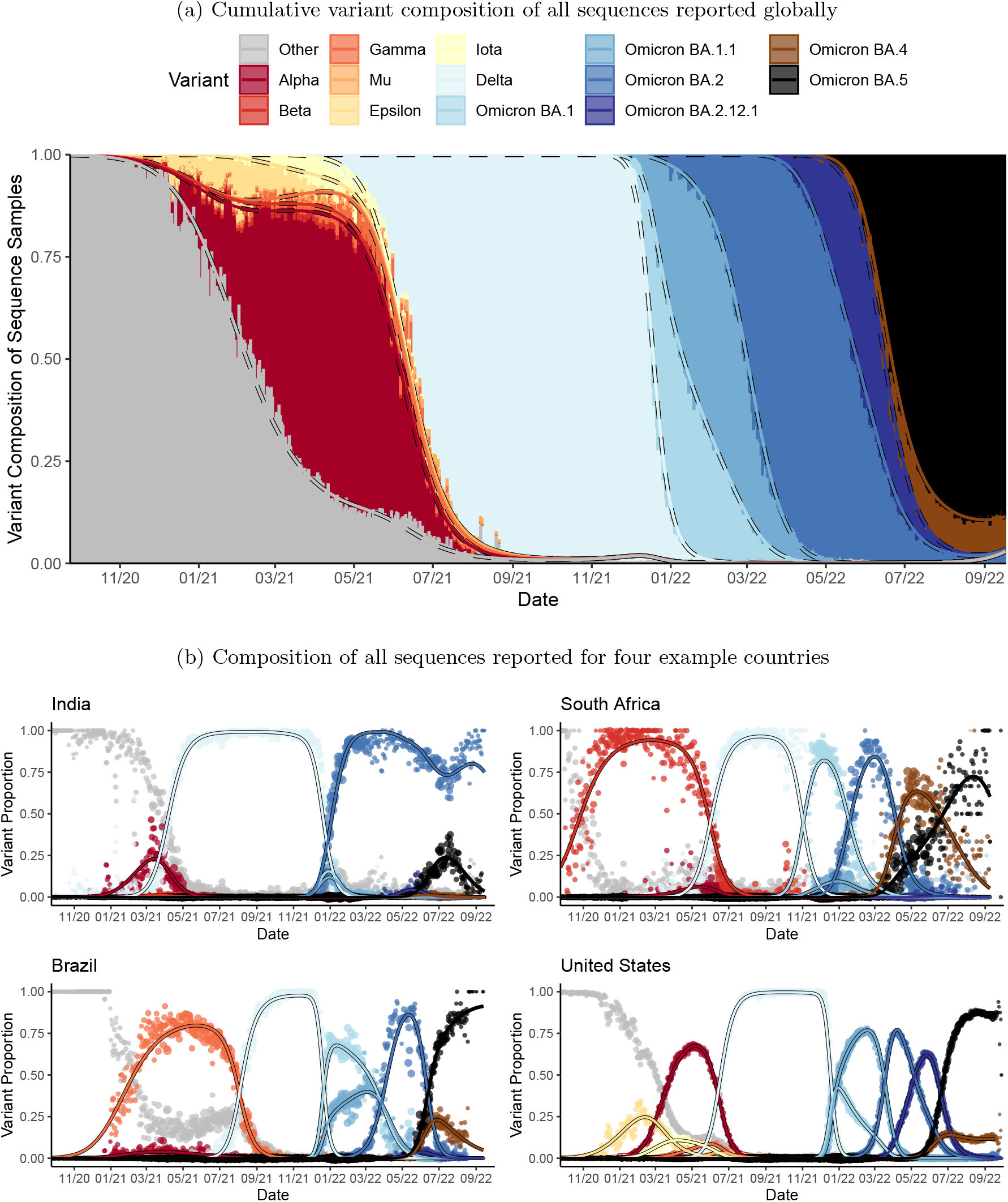
Daily variant composition of all SARS-CoV-2 sequences reported to GISAID globally and for four example countries (points) along with fitted variant proportions (lines) from the primary analysis. Fitted lines show the point estimates obtained from fitting the multinomial model in *Eq. 1*. The size of the plotted points correspond to the total number of sequenced samples, relative to the daily maximum within each country. Variants in the emerging variants sub-analysis are not shown separately.

#### Clinical and demographic data

Daily confirmed COVID-19 case and death data were obtained from the Johns Hopkins Center for Systems Science and Engineering (CSSEGIS), along with daily Oxford COVID-19 Government Response indicator (0=none, 100=strict) and WorldPop age, population density, and population information for each location.^2,14,15^ Daily model-predicted mask usage (%) based on survey data was obtained from the Institute for Health Metrics and Evaluation (IHME) at the University of Washington.^16^ Population percent with less than secondary education and the average disposable income (in dollars) were obtained from the Organization for Economic Co-operation and Development (OECD).^17^ When missing, region-level OECD data were assigned the reported country-level value. Additional information is provided in **Supp. Table A.1**.

### 2.2 Characterizing speed, timing, and magnitude of SARS-CoV-2 variant transitions across locations and variant categories

The variant landscape in a given population is dynamic, with the number of competing variants changing through time (**Figure 1**). We propose a model for the variant proportions over time for each location that directly accounts for multiple competing variants. Several similar (but often less flexible) models of SARS-CoV-2 variant transitions have been proposed elsewhere.^18,19,20,21,22,23^ Depending on data availability, our primary analysis considered up to 13 variant Pango lineage categories for each location, including Alpha, Beta, Iota, Gamma, Mu, Epsilon, Delta, Omicron BA.1 (excluding BA.1.1), Omicron BA.1.1, Omicron BA.2 (excluding BA.2.12.1), Omicron BA.2.12.1, Omicron BA.4, and Omicron BA.5, as well as “others”.

Let *y_ij_*(*t*) be the observed number of sequences for location *i* and day *t* attributed to variant/sub-variant *j*, and let *y*_*i*0_(*t*) represent the number of sequences in the “other” category. For each day, we defined the true *proportion* of sequences attributed to variant *j* as *p_ij_*(*t*), with *p*_*i*0_(*t*) representing “other.” We assumed an independent multinomial distribution for variant composition of sequencing, with proportions modeled as:

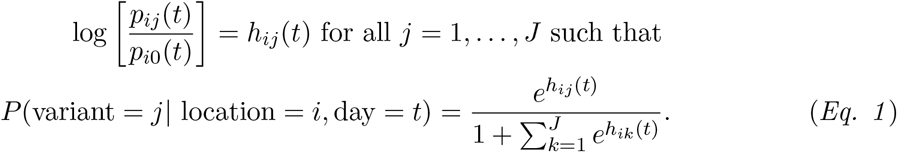

This model is similar to one in Figgins and Bedford^22^, who noted convenient parameter interpretations but poor data fits when *h_ij_*(*t*) is linear in *t*. We posited a more flexible cubic spline model for each *h_ij_*(*t*), with a knot at the median of *t*. Multiple knots and additional linear terms did not further improve the results. Resulting fitted variant proportions are illustrated in **Figure 1**.

Noting that 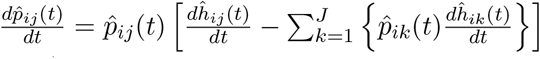, we summarized fitted 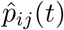 using the following metrics:

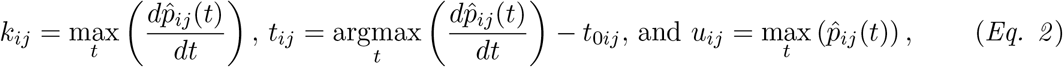

where *k_ij_* represents the maximum slope of the variant transition curve (i.e., transition speed), *t_ij_* is the relative timing in days at which the maximum slope is achieved (i.e., transition timing), and *u_ij_* is the maximum fitted variant prevalence (i.e, transition peak prevalence). The earliest transition time is set to zero for each variant *j*, with 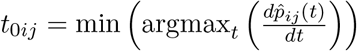. Higher *k_ij_* and smaller *t_ij_* corresponds to steeper and earlier variant transitions, respectively.

**Supp. Figure A.2** provides our criteria for determining which location and variant combinations were modeled. After fitting *Eq. 1*, we excluded some locations based on visual evaluation of the fits. Out of 590 country/regions considered, 215 locations (and a total of 1,580 variant transitions) were included in the primary analysis (**Supp. Figure A.4**).

Emergent Omicron Pango lineage groupings were not considered in our primary analysis due to insufficient data. In a special emerging variants sub-analysis, we fit *Eq. 1* and calculated summary metrics to characterize currently-available data for the Omicron BA.2.75, XBB/XBB.1, and BQ.1 variants; these estimates may change substantially as more data are collected. Unless otherwise stated, all analyses used Alpha through Omicron BA.5 variant groupings.

### 2.3 Clustering locations in terms of similarity in SARS-CoV-2 variant transition profiles

To characterize location similarities across Omicron transitions, we performed a hierarchical clustering analysis. Since the included sub-variants differed by location, we again summarized *Eq. 2* metrics by fitting the following regression model:

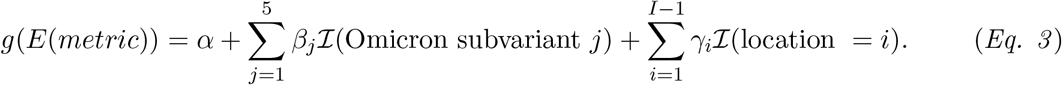

Estimated *γ_i_*’s capture the average difference between each location and the reference location (USA, due to large sample size) in terms of the summary metric across Omicron transitions. We chose Gaussian (log_10_(*k*)), Poisson (*t*), and Beta (*u*) regressions using canonical link functions *g*. Clustering was performed only for 155 locations with at least 4 included Omicron sub-variants. Emerging Omicron sub-variants were not separated out in this analysis.

We then performed a Wald agglomerative hierarchical clustering on *γ* estimates from *Eq. 3* using the R package *cluster*^24^. In defining the number of clusters, we compared cluster size, within cluster sum of squares, intra-cluster variance, and how South Africa was clustered (since its transition dynamics were distinctive). **Supp. Figure B.1** illustrates the resulting 7 clusters in terms of their *Eq. 3* coefficients.

### 2.4 Exploring relationships between variant transition metrics and contemporaneous disease landscape

We obtained location attributes at the time each 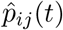 first reached 5%. We chose 5% to focus on the critical time when the new variant is gaining a foothold locally but clinical surveillance would likely not have been appreciably impacted. Characteristics of interest included demographics (e.g., population density), COVID clinical landscape (e.g., case burden and vaccination rates), and current COVID-related public policy (e.g., Governmental response indicator). We also identified two proxies for existing variant competition at the time of the new variant emergence in each location: 1) the number of co-circulating SARS-CoV-2 variants/sub-variants with at least 5% prevalence and 2) the competition ratio, defined as the maximum percent increase in existing variants’ prevalences between 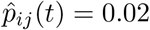 and 0.05.^25^ Additionally, we included the timing and height of the most recent prior COVID-19 case wave peak (**Supp. Section A**).

For each variant, we calculated the Spearman correlation between the *Eq. 2* metrics and location characteristics along with cross-correlations of transition summary metrics across variant waves. For each summary metric, we then performed random forest modeling to study the adjusted location-transition associations. We used the R package *randomForest*^26^, with missing predictor data handled using proximity-based imputation. Out-of-bag importance metrics were calculated based on 10,000 regression trees. We also fit regression models for each of log_10_(*k*), *t*, and *u*, using Gaussian, Poisson, and Beta regression, respectively. Prior to regression modeling, missing data were handled by multiple imputation using the R package *mice*^27^. All models were also adjusted for variant/sub-variant. **Supp. Figures A.3** and **D.1** describe the data missingness and model goodness of fit. Modeling results based only on locations with complete data are provided in **Supp. Figure D.2**.

## 3 Results

### 3.1 Characterizing speed, timing, and magnitude of SARS-CoV-2 variant transition profiles across locations and Pango lineage groups

**Figure 2** summarizes the fitted variant transitions from the primary analysis, along with three emerging Omicron sub-variants. **Supp. Figures C.1-C.3** map estimates for several variants of interest. The Beta, Gamma, Mu, Epsilon, and Iota variants were associated with lower variant prevalence (u) and transition speeds (*k*), except for the Gamma transition in South America and the Beta transition in Southern Africa. Delta and Omicron BA.1, BA.1.1, BA.2, and BA.5 variants tended to have fast (high *k*) transitions, although there was substantial variability in terms of speed and prevalence attained by Omicron BA.1 and BA.1.1 globally. The Omicron BA.1.1 variant achieved a strong presence in the Americas, reaching a prevalence of about 75% in the USA, where it had a relatively early start (**Figure 1b**). Alpha had a slow and small transition in South America, likely due to competition with Gamma and Mu, and in South Africa, where it was competing with Beta. In contrast, the transition speed and maximum prevalence had little heterogeneity for the Delta, exhibiting a rapid and total transition in most locations. Omicron BA.4 and BA.5 were first observed in South Africa and spread globally at roughly the same time (**Figure 2**); however these lineages had profoundly different trajectories in terms of their maximum transition slopes, maximum prevalence, and their relative time to transition, suggesting selective advantage of BA.5 over BA.4 (**Figure 2**). Of note, the founder forms and early expansions of BA.4 and BA.5 carried identical Spike sequences,^28^ implicating changes outside of the spike protein in the observed differences. All three newly emerging Omicron variants (BA.2.75, XBB/XBB.1, and BQ.1) had transition slopes on par with the earlier Omicron sub-variants. Among the small number of locations that had reached their maximum transition slope by our last day of sampling (11/03/2022), XBB/XBB.1 and BQ.1 generally had higher transition slopes than BA.2.75.

**Figure 2:**
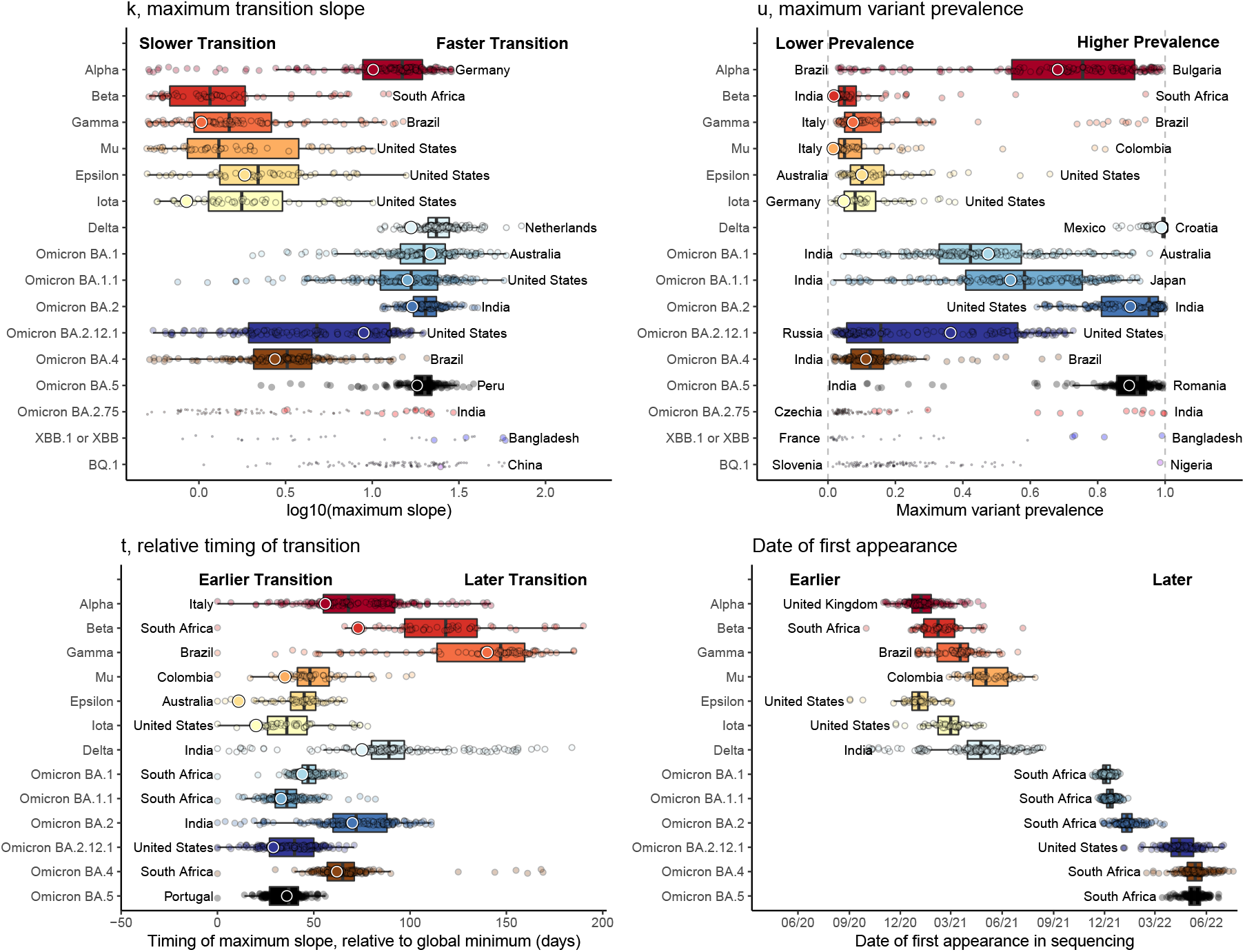
Distribution of highest variant transition speeds (*k*), relative timings (*t*), and highest variant prevalences (*u*) across 215 locations, with text annotations indicating the countries having the highest and/or lowest value. Large circles correspond to global estimates based on analyzing all locations together. For emerging variants Omicron BA.2.75, XBB.1/XBB, and BQ.1, medium- and small-sized circles provide estimates for locations that have and have not reached their maximum fitted slope by 11/03/2022, respectively. For these transitions, maximum slope estimates are expected to *substantially increase* as more data become available. Many maximum variant prevalences are also likely to increase as more data are collected for all locations. Small global estimates of k for Mu (0.34) and Beta (0.26) variants are not shown. These results demonstrate substantial heterogeneity in variant transition dynamics between locations.

The date of “first” appearance (i.e., the first day with at least two sequences) provides insight into the relative timing of each variant’s global spread. Some variants (e.g., Beta and Epsilon) were first sequenced in the originating country long before they were more commonly sequenced globally. In contrast, the Omicron sub-variants were observed globally very quickly after their discovery.

The relative timing of the maximum transition slope, *t*, is defined in terms of days since the earliest global transition for each variant. This metric is distinct from the first variant appearance, since a variant can circulate at low levels for a long time before gaining a foothold in a given location. Therefore, *t* provides a better metric for the variant transition *timing*. The Beta and Delta waves hit much earlier in their originating countries (South Africa and India, respectively) than they did globally, with some countries’ Delta variant transition occurring over 6 months later. In contrast, the Omicron BA.1, BA.1.1, and BA.5 waves occurred more quickly and with much less variability globally. Low transition timings for Mu, Epsilon, and Iota are due to limited localized spread.

### 3.2 Clustering locations in terms of similarity in SARS-CoV-2 variant transition profiles

To evaluate whether groups of geographic locations tended to have shared patterns *across* Omicron transitions (in the primary analysis), we performed hierarchical clustering, using data through September, 2022. The resulting seven clusters are illustrated in **Figure 3**.

**Figure 3:**
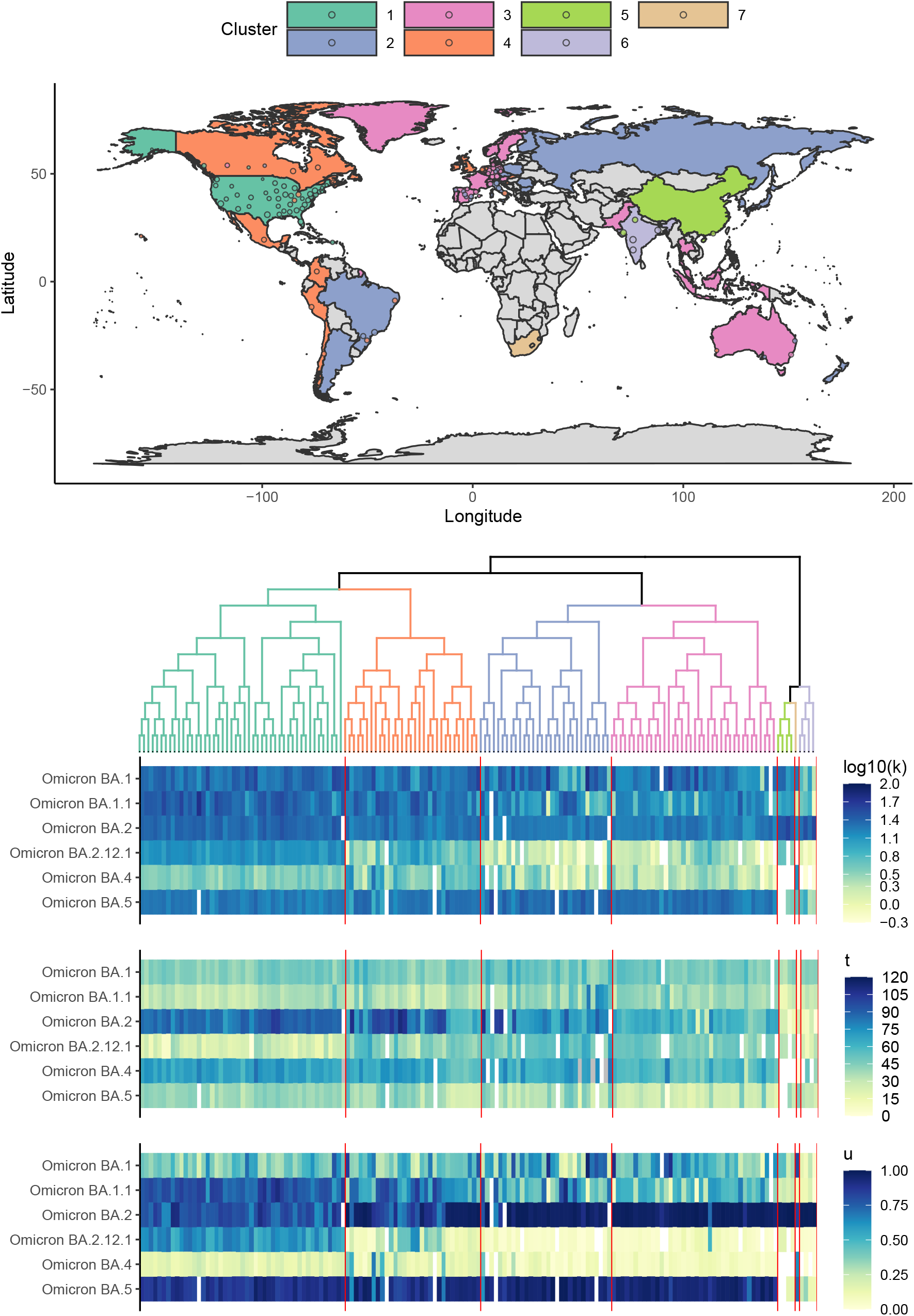
Hierarchical clustering of *k* (maximum transition slope), *t* (relative timing of transition), and *u* (maximum prevalence) across all Omicron variant waves included in the primary analysis. The semi-transparent circles overlaid on the map provide estimates for included sub-national region locations. Some sub-national regions outside of contiguous national boundaries (e.g., Greenland, a sub-region of Denmark) are instead filled in with the appropriate color to reflect the regional value. Countries shown in grey are those for which data were either unavailable or insufficient. The heatmap illustrates the estimated summary metrics for all locations and all Omicron variant transitions considered. South Africa, India, and China were notable for their distinctive transition dynamics.

The first cluster (mostly the United States) was distinctive due to its pronounced and early BA.2.12.1 transition and substantial BA.1.1 transition. The second (Eastern Europe, Russia, and part of South America), third (primarily Western Europe and Australia) and fourth (The United Kingdom, Mexico, Canada, and part of South America) clusters tended to be comparatively similar on average, with the fourth cluster having slightly higher BA.2.12.1 prevalences on average. The fifth and sixth clusters, consisting of China and India, were distinctive in that the Omicron transitions were dominated by Omicron BA.2 only, with comparatively low prevalence of all other variants, including Omicron BA.5. BA.5 had just begun to increase in India when the BA.2 sublineage BA.2.75 rapidly became the dominant form regionally (**Figures 1 and 2**). The seventh South Africa cluster led the world in earliest Omicron transitions. Omicron BA.1 and BA.4 transitions were particularly rapid and large in South Africa, which had a comparatively low rate of BA.1.1 expansion despite its earliest detection there. As illustrated in **Supp. Figure B.1**, the China, India, and South Africa clusters were clear outliers.

### 3.3 Exploring relationships between variant transitions and contemporaneous disease landscape

The clustering analysis indicated that Omicron variant transitions tended to be more similar between some location pairs than others, suggesting there may be a link between transition dynamics and location characteristics. In **Supp. Figure C.6**, we explored correlations between transition summary metrics and location characteristics (**Supp. Figure C.6**).

In **Figure 4**, we investigate the relationship between the maximum transition slope and the number of co-circulating variants when each variant reached 5% prevalence. A higher number of co-circulating variants was significantly associated with lower transition speeds for many variants after multiple testing adjustment, including Delta, Epsilon, Gamma, and Omicron BA.1, BA.2, and BA.5.

**Figure 4:**
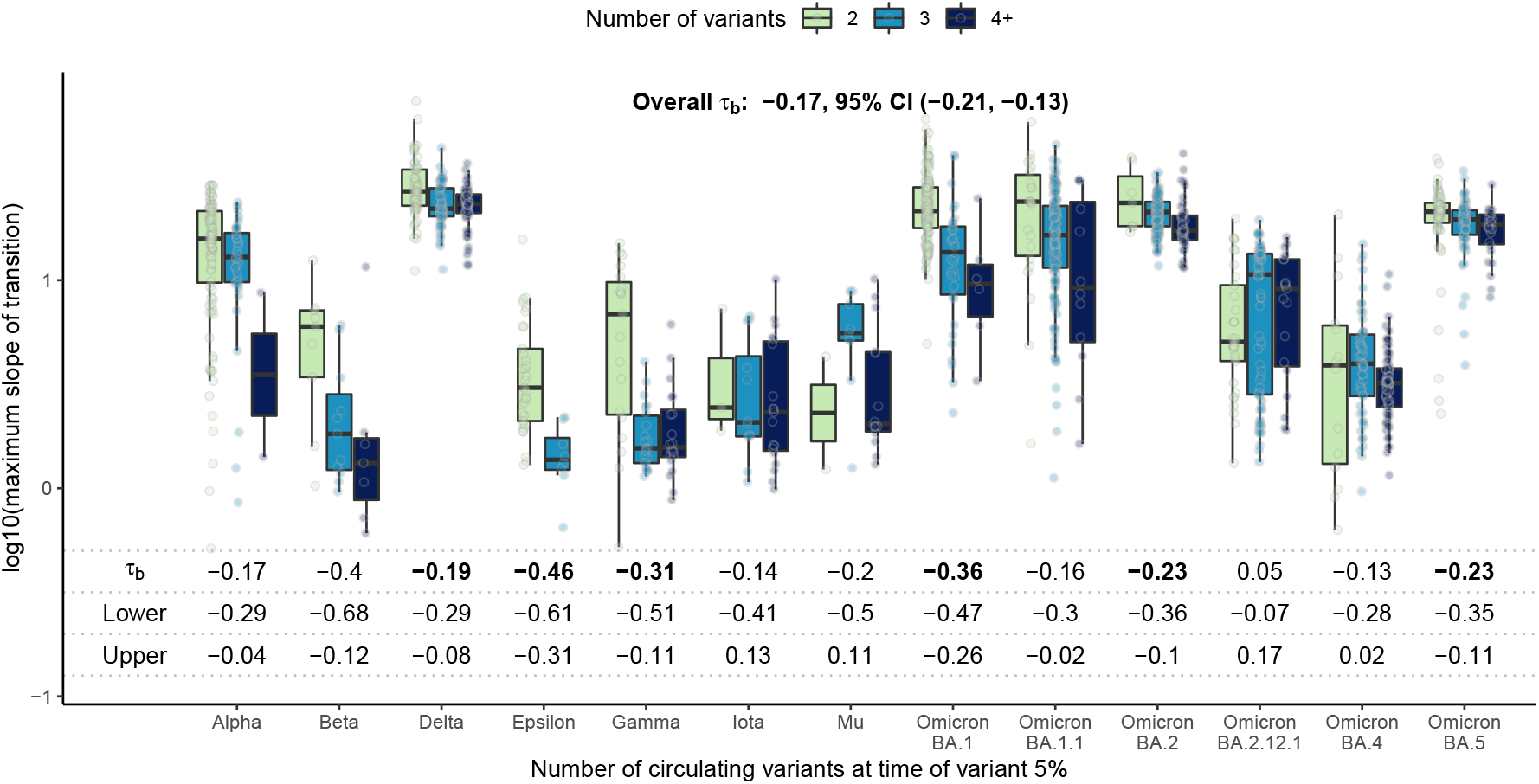
Global estimates of k, the maximum slope of the variant transition curve, by the number of co-circulating variants (including the variant itself) at the time of variant 5% prevalence. Kendall’s *τ_b_* correlation and corresponding 95% confidence intervals are also provided. Bolded correlation estimates are significantly non-zero, after adjusting for multiple testing using a Bonferroni correction (14 tests). A higher number of co-circulating variants was significantly associated with lower transition speed for many variants.

In **Figure 5**, we plot variant transition summary metrics as a function of population vaccination rates. **Supp. Figure C.7** provides correlation estimates by variant. Higher vaccination rates were associated with later and slower global spread before the Mu and Delta variants emerged, when vaccination rates were generally low. For Omicron, however, vaccination rates were at most weakly associated with the speed and timing of variant transitions.

**Figure 5:**
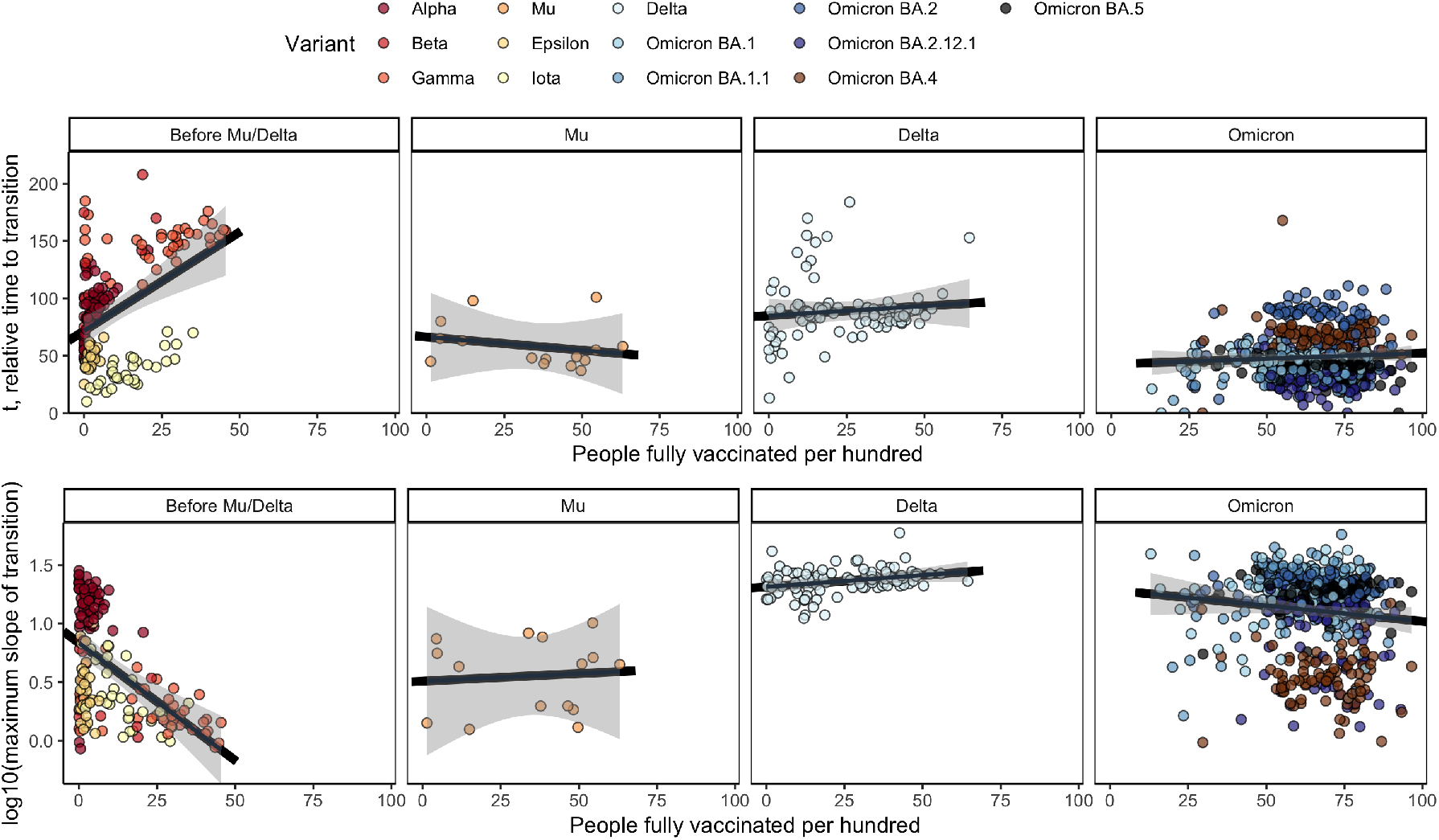
Global estimates of *t* and *k*, the timing and magnitude of the maximum slope of the transition curve, by the vaccination rate at the time of variant 5%prevalence. 95% confidence bands (multiple testing adjusted) for the fitted linear regression for each panel are shown in gray. Higher vaccination rates were significantly associated higher transition speed and later transitions prior to the Mu/Delta variants. We did not observe a significant association between vaccination and transition properties for Omicron sub-variants, after correcting for multiple testing.

We then estimated the *adjusted* associations between location characteristics and the transition summary metrics using both random forest and regression modeling (**Figure 6**). We used two modeling approaches, since each contributes a different element of the story. Random forest modeling accounts for complicated interactions between variables, while regression provides interpretable parameter estimates. All models also adjusted for variant, which was generally the most important predictor of each summary metric (not shown).

**Figure 6:**
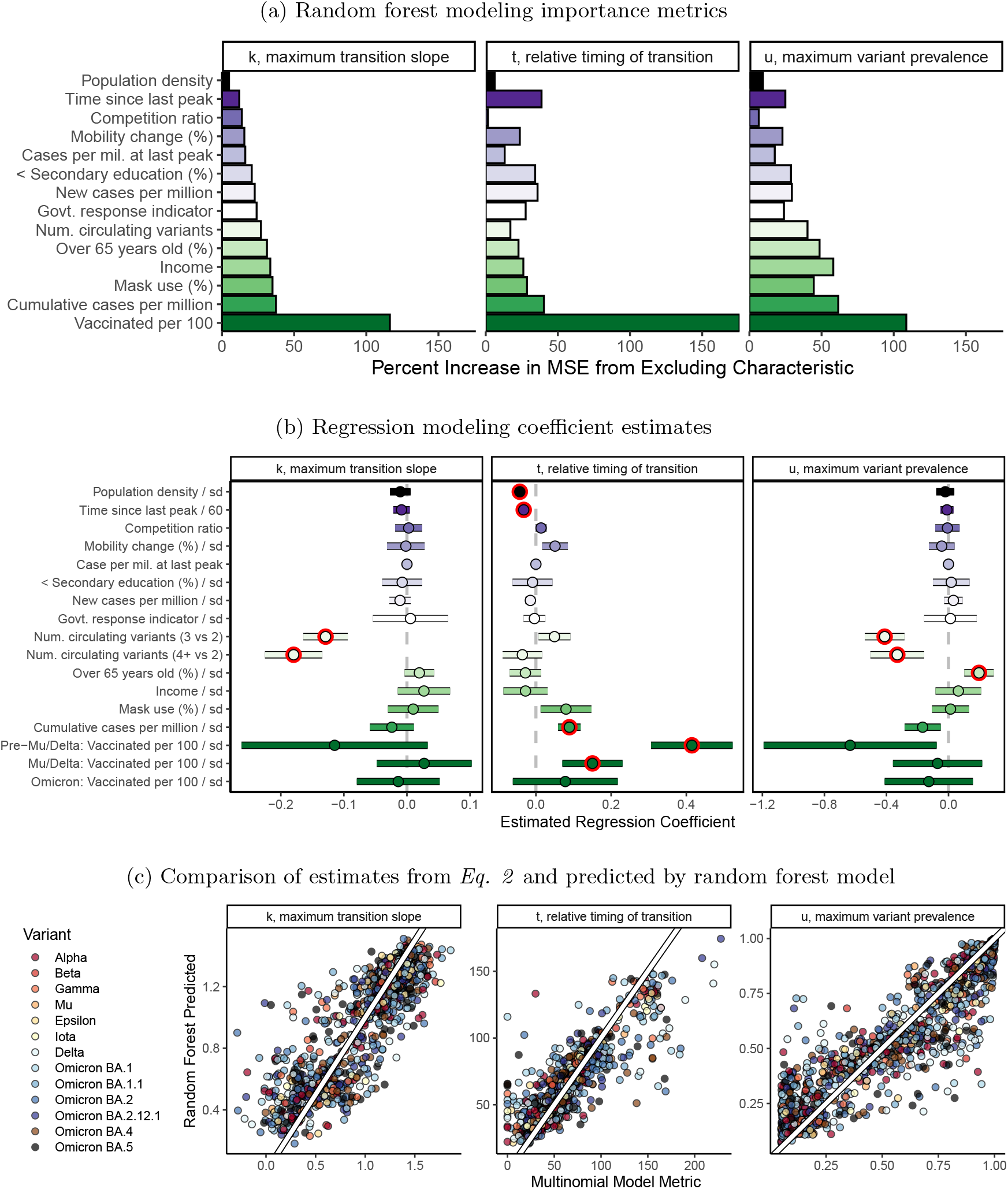
Relative importance (a) and regression model coefficient estimates (b) of two adjusted models for associations between location characteristics and variant transition summaries. A comparison of random forest-predicted summary metrics to estimates from *Eq. 2* is shown in (c). Red circles in (b) denote associations statistically significant at the 0.05 level after Bonferroni multiple testing correction. Vaccination rates, cumulative prior cases per million, population density, population age, the time since the last case wave peak, and the number of co-circulating variants were all significantly associated with variant transition dynamics after adjusting for other location characteristics and accounting for multiple testing.^1^

Even after adjusting for location characteristics and multiple testing, vaccination status was significantly associated with variant transition dynamics pre-Mu/Delta. In particular, one standard deviation higher vaccination rate per 100 was associated with a 51% (95% CI: 36-69%) *later* time to variant transition. Higher vaccination rates per 100 were also associated with lower transition peak prevalences pre-Mu/Delta. These strong associations were not observed during the Omicron waves and were not observed or were attenuated during the Mu and Delta waves.

A higher number of co-circulating variants was strongly associated with slower and lower peak-prevalence variant transitions (p-value < 0.0001 for each). A higher prior COVID-19 case rate per million, a shorter time since the last case wave peak, and lower population density were all significantly associated with *later* variant transitions. Locations with higher proportions of the population over 65 tended to have higher peak variant prevalences.

## 4 Discussion

Although highly efficacious COVID-19 vaccines were developed with unprecedented speed and have substantially helped temper the impact of the pandemic, the continuing evolution of SARS-CoV-2 has been associated with new waves of disease spread and with viral variants that have become progressively more infectious and resistant to protective antibodies.^7,8,11^ A variant with a selective advantage can quickly become the most prevalent viral after its emergence and can radically change the clinical landscape of virus transmission. An emerging variant’s relative selective advantage is reflected in the speed, timing, and magnitude with which the emerging variant out-competes existing variants in a given country. In this work, we characterized variant transitions across 13 SARS-CoV-2 variants and 215 countries and sub-country regions throughout the COVID-19 pandemic. We also explored relationships between properties of the variant transitions and contemporaneous disease landscape (e.g. vaccination rates, natural immunity due to past infections, and demographics). Through an emergent variant sub-analysis, we also illustrated how these metrics can be used to monitor ongoing variant transitions. In this early analysis, we found that the transitions to BQ.1 and XBB in countries where they have already become established was relatively rapid when compared to prior Omicron variant transitions (**Figure 2**). Although this analysis is preliminary and by necessity based on a small number of countries, it is consistent with a selective advantage, providing further impetus for current efforts to better resolve biological characteristics of these viral lineages.

In this paper, we demonstrated that historical variant transition dynamics differed substantially between locations (**Figure 2**) and were associated with vaccination rates, prior infection rates, the time since the last COVID-19 peak, population demographics, and the number of co-circulating variants in competition with the emergent variant (**Figure 4–6**). In particular, stronger natural immunity (due to higher prior infection rates and a shorter time since the last COVID-19 peak) was strongly associated with later variant transitions relative to other countries, suggesting that the new variant transitions may be slower in locations with a large recent disease burden, consistent with protective antibodies being at higher levels due to recent stimulation, and potentially being more cross-reactive if they were elicited by a variant that was more closely related to a newly emergent form.

The association between vaccination rates is particularly interesting; while higher vaccination rates were associated with slower transitions prior to Delta and Mu, the Delta and Mu variants were key inflection points. Among Omicron variants, the association was strongly attenuated (**Figure 5**), consistent with Omicron’s resistance to vaccine-elicited neutralizing antibodies, which are a key aspect of protection from infection.^7,8,11^ The ability of bivalent vaccines to offer additional protection against Omicron-related infections is still being resolved,^29,30^ and the neutralizing antibody sensitivity of emergent variants may impact the ability of vaccine boosters to slow transition times to new variants going forward.

The analyses in this paper are subject to many potential biases. Firstly, the SARS-CoV-2 sequences that are available may not be representative of circulating variants. For example, sequencing efforts may over-sample a large outbreak or over-sample cases tied to an emerging variant. Future work should explore strategies for quantifying biases in GISAID sequence reporting by location over time. To add additional complication, data quality issues such as strings of ambiguous base calls can result in Pango designation mis-assignments, and changing Pango lineage designations as the virus evolves can obfuscate emerging variant transitions. Confirmed COVID-19 case and vaccination data are also imperfect, with substantial under-reporting that likely varies over time. Missing data also presents a challenge, and the imputation methods we have used to address the missing data have implicit assumptions about the representativeness of the observed data.

Although there has been a remarkable global effort to track and understand transitions during the pandemic, still SARS-CoV-2 and COVID-19 data streams are biased toward higher-income countries, since low- and middle-income countries tend to have less complete data and to submit fewer sequences to GISAID. Because we had inclusion requirements based on completeness and volume of sequence submissions, many variant-by-location combinations were omitted. As a result, our analyses are implicitly biased toward data collected in higher-income countries, as shown in **Supp. Figure A.4a**.

Overall, this analysis highlights the complicated relationships between variant transitions and the contemporaneous immunologic and clinical context. Additionally, our results demonstrate substantial heterogeneity in how an emerging variant interacts with co-circulating variants across locations. Future work may be able to leverage this heterogeneity and data on historical variant transitions to help forecast how emergent variants may behave in the future, potentially using observed transitions in the variant’s country of origin to forecast the variant’s future transition properties in other countries.

## Acknowledgments

This work was partially funded by the Laboratory Directed Research and Development (LDRD) Exploratory Research Project 20220660ER. Dr. Beesley was funded by the LDRD Richard Feynman Postdoctoral Fellowship 20210761PRD1. Dr. Wagh also supported by LDRD 20220399ER. This work is approved for distribution under **LA-UR-22-32123**. The findings and conclusions in this report are those of the authors and do not necessarily represent the official position of Los Alamos National Laboratory. We gratefully acknowledge all data contributors (i.e., the authors and their originating laboratories responsible for obtaining the GISAID data specimens) and their submitting laboratories for generating the genetic sequence and metadata and for sharing via the GISAID Initiative, on which this research is based.

## Data Sharing Statement

All data used in this analysis are publicly available. Details are provided in **Supp. Table A.1**.

## Footnotes

1 Random forest importance measured in terms of percent increase in mean squared prediction error. For regression modeling, continuous predictors were scaled by their standard deviations. Gaussian, Poisson, and Beta regression were used for log_10_(*k*), *t*, and *u*, respectively. All models also adjusted for variant/sub-variant. Missing predictor information was handled using imputation.

