## Supplementary Materials for "SARS-CoV-2 variant transition dynamics are associated with vaccination rates, number of co-circulating variants, and natural immunity"

November 18, 2022

#### A Data sources and processing

In this section, we provide additional information about the data sources and data resolutions used. **Figure A.1** illustrates the data sources used in this work along with the data spatial and temporal resolutions. **Table A.1** provides accession information for the data.

Sub-national regional data (as defined by the Johns Hopkins CSSEGIS) were used for Australia, Brazil, Canada, Chile, China, Colombia, Germany, India, Italy, Mexico, the Netherlands, Peru, Russia, Spain, and the United States[1]. **Figure A.2** describes the total number of countries/sub-national regions included in the final analysis, and **Figure A.4** provides the number of locations (countries/regions) included in the analysis of each SARS-CoV-2 variant considered. Definitions for each variant based on the pangolineage designations are provided in **Table A.1**.

Case peaks were identified using a finding algorithm designed to capture true case peaks while ignoring minor oscillations in the face of highly variable data quality and noise level. The input to the algorithm is the set of all (date, number of cases) observations for a given location, i.e. a time series of case counts. For each location, we first smooth the time series via LOESS to create a smoothed case profile. Then, we find peaks (and troughs) of the smoothed case profile by identifying locations where the rolling max (min) occurs at its own location. We correct the identified peaks (troughs) found originally using the smoothed case profile by checking to see if any of the closest neighboring dates have a higher (lower) case count. We require unique, alternating peaks and troughs, prioritizing larger peaks and smaller troughs. Given this alternating set of peaks and troughs, we use % increase since last trough and distance to former peak to remove identified peaks that represent oscillations rather than visually significant/distinct peaks. We perform further manual processing after visually inspecting the identified peaks and troughs overlaid on the time series; we then rerun the algorithm with less smoothing (i.e., span set to 0.075) for countries with close-together peaks that weren't captured due to over-smoothing. This semi-automated, semi-manual process is implemented for each location considered to identify a set of dates corresponding to peaks in the confirmed case count data.

Figure A.1: Illustration of data sources and their temporal and spatial resolutions

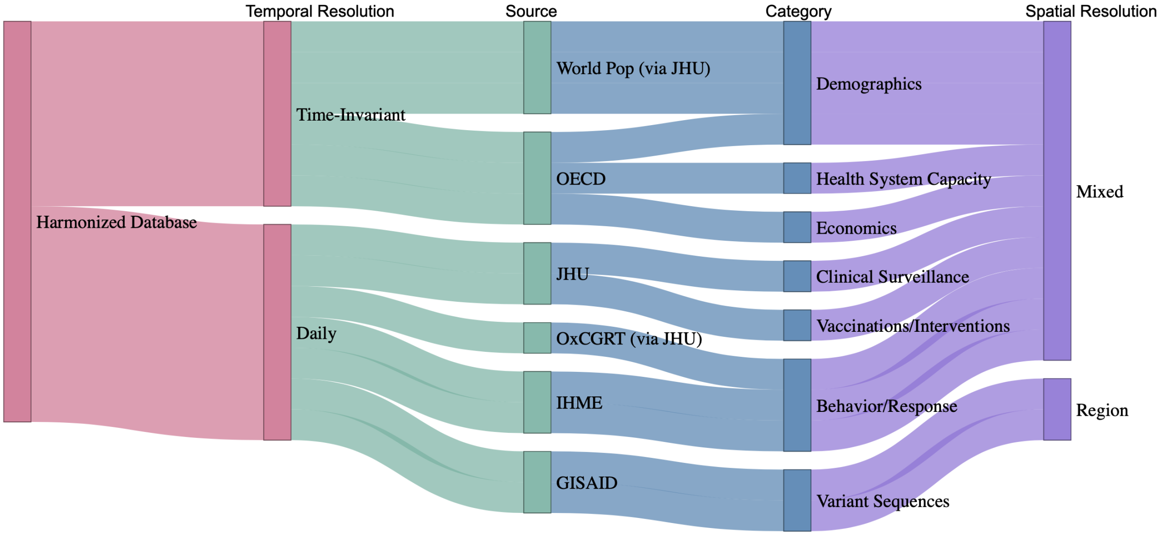


Table A.1: Data sources and accession information for data used in primary analysis. Time coverage corresponds to the longest time period with available data for any country. "Pre-pandemic" indicates a time-invariant variable collected for some pre-pandemic time. All data in the primary analysis were collected on 10-01-2022 or 10-03-2022 except for OECD, which was collected on 03-21-2022. The emerging variants analysis included GISAID data through 11-03-2022.

| **Data Source** | **Variables** | **Time Coverage**[1] | **Reference** |
|---|---|---|---|
| Johns Hopkins CSSEGIS | Cases<br>Vaccinations | 02/2020 to 10/2022<br>12/2020 to 10/2022 | Badr et al. [1] |
| Oxford COVID-19 Government Response Tracker (OxCGRT) via JHU | Governmental response indices | 01/2020 to 10/2022 | Thomas et al. [2] |
| WorldPop via JHU | Median age<br>Population density<br>Population | Pre-pandemic<br>Pre-pandemic<br>Pre-pandemic | WorldPop [3] |
| GISAID | SARS-CoV-2 sequences<br>Sequence date/location/Pango lineage | 12/2019 to 10/2022 | Khare et al. [4] |
| Institute for Health Metrics and Evaluation (IHME) | Model-based mask use | 02/2020 to 10/2022 | IHME [5] |
| Organisation for Economic Co-operation and Development (OECD) | Education (% less than secondary)<br>Disposable income | Pre-pandemic<br>Pre-pandemic | OECD [6] |

Table A.2: Variant definitions as a function of Pango lineage

| **Variant/sub-variant** | **Pango lineage** |
|---|---|
| *Primary Analysis* | |
| Alpha | B.1.1.7 or B.1.1.7.* or Q.* |
| Beta | B.1.351 or B.1.351.* |
| Gamma | B.1.1.28 or B.1.1.28.* or P.1 or P.1.* |
| Mu | B.1.621 or B.1.621.* or BB.2 |
| Epsilon | B.1.427 or B.1.427.* or B.1.429 or B.1.429.* |
| Iota | B.1.526 or B.1.526.* |
| Delta | B.1.617.2 or B.1.617.2* or AY.* |
| Omicron BA.1 (except BA.1.1) | BA.1 or BA.1.* or BD.*, excluding BA.1.1, BA.1.1.*, and BC.* |
| Omicron BA.1.1 | BA.1.1 or BA.1.1.* or BC.* |
| Omicron BA.2 (except BA.2.12.1) | BA.2 or BA.2.* or BS.* or BH.* or BP.* or BJ.* or BL.* or CA.* or BM.* or BR.* or BN.* or BY.* or CB.*, excluding BA.2.12.1, BA.2.12.1.*, and BG.* |
| Omicron BA.2.12.1 | BA.2.12.1 or BA.2.12.1.* or BG.* |
| Omicron BA.4 | BA.4 or BA.4.* |
| Omicron BA.5 | BA.5 or BA.5.* or BE.* or BF.* or BK.* or BQ.* or BT.* or BU.* or BV.* or BW.* or BZ.* or CC.* or CF.* or CD.* or CE.* |
| *Emerging Variants Analysis* | |
| Omicron BA.2.75 | BA.2.75 or BA.2.75.* or BL.* or CA.* or BM.* or BR.* or BN.* or BY.* or CB.* |
| XBB.1 or XBB | XBB.1 or XBB.1.* or XBB |
| BQ.1 | BQ.1 or BQ.1.* or CZ. or CW. |

[1] 'XX.*' indicates all sequences starting with the characters 'XX.'

Figure A.2: Visualization of inclusion and exclusion criteria to obtain analytical sample for the primary analysis. Numbers following each country indicate the number of sub-country regions included. Exclusion criteria were weakened for emerging variants analysis, where reporting gaps up to 5 days (rather than 3) were allowed and we required at least 5 sequences reported for at least 30% of days (rather than 50%). The emerging variants analysis included 84, 29, and 100 locations for BA.2.75, XBB.1/XBB, and BQ.1, respectively.

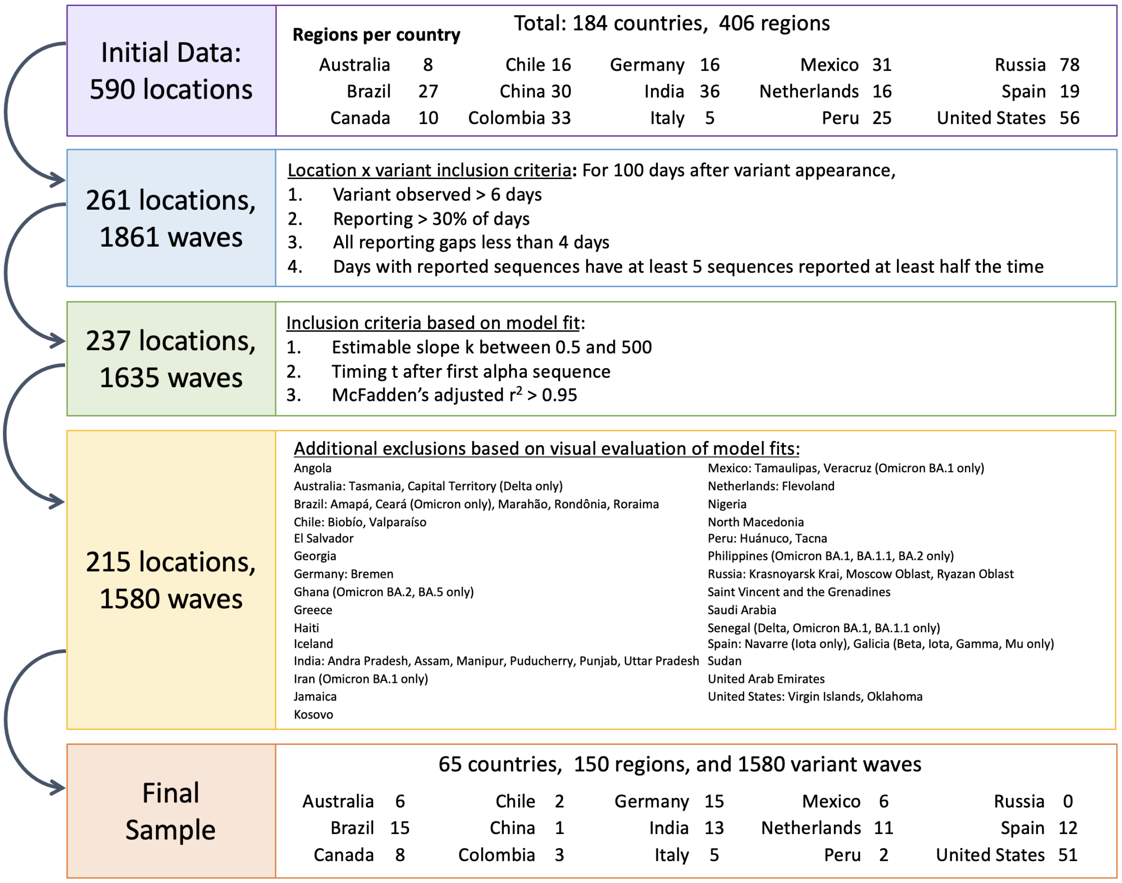


Figure A.3: Percent of included locations with observed data by variant for each location characteristic considered in primary analysis.

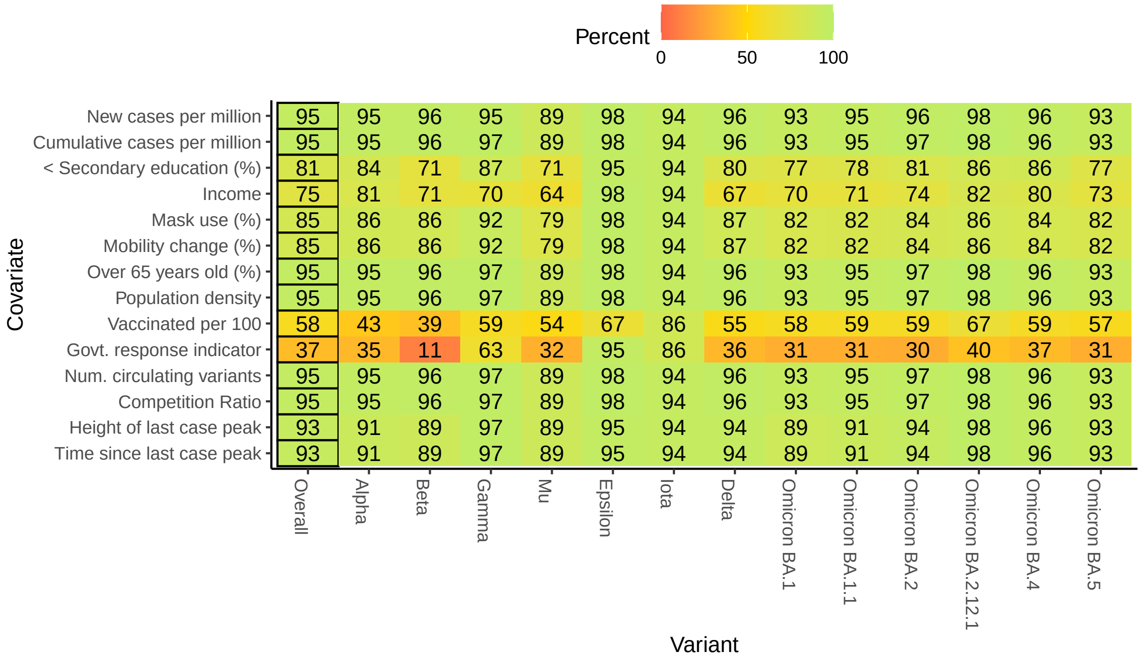

Figure A.4: Locations included in the primary analysis among the initial 590.[1]

(a) Map of included and excluded locations

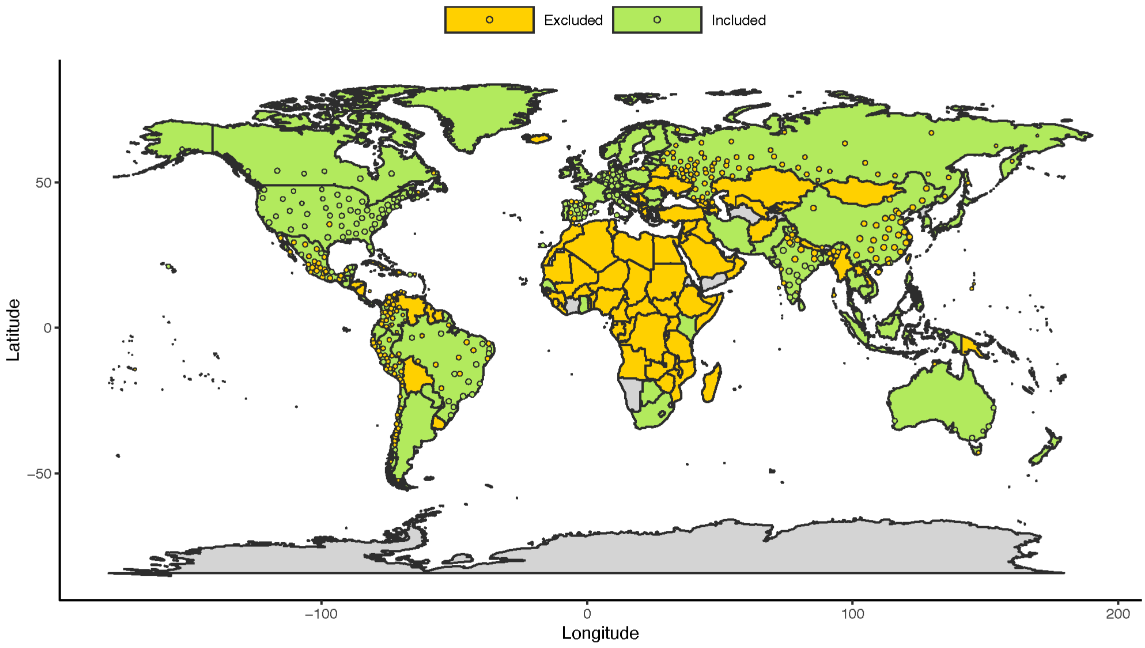


(b) Number of included locations by variant

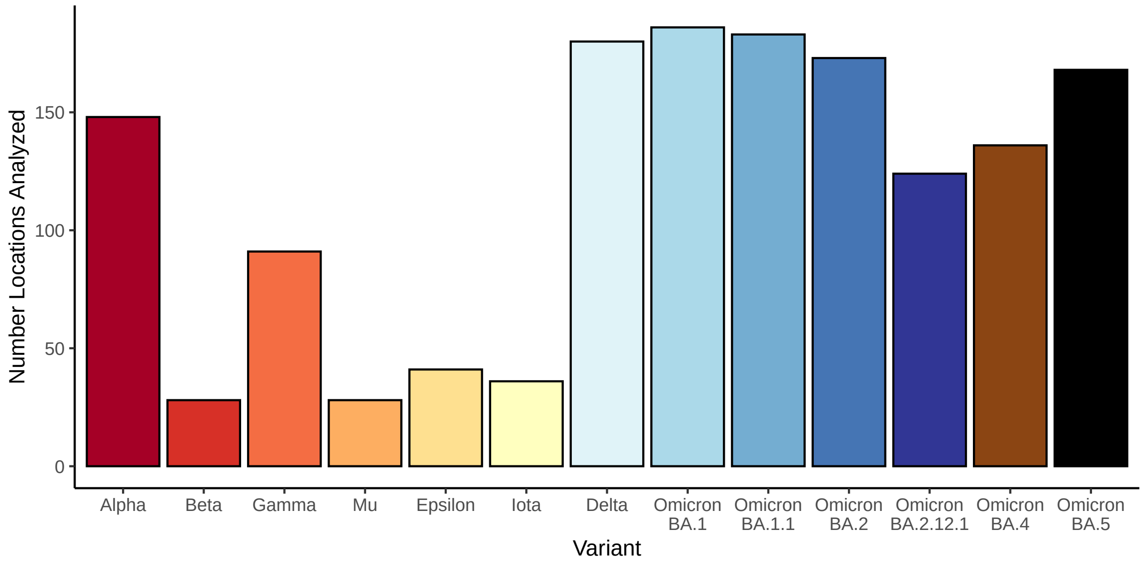


[1] Locations were excluded due to the frequency and volume of sequence data reporting, along with the goodness of fit of the multinomial regression modeling.

### B Hierarchical clustering of variant transition summary estimates

Figure B.1: Clusters identified by hierarchical clustering of all $k$, $t$, and $u$ estimates for each location. Locations included in each cluster are shown in **Figure 3**. Clustering was based on variant designations in primary analysis (rather than the emerging variants sub-analysis). In each panel, the plotted x-y location of each point corresponds to the location's fitted coefficients $\gamma$ from *Eq. 3* for different summary metrics.

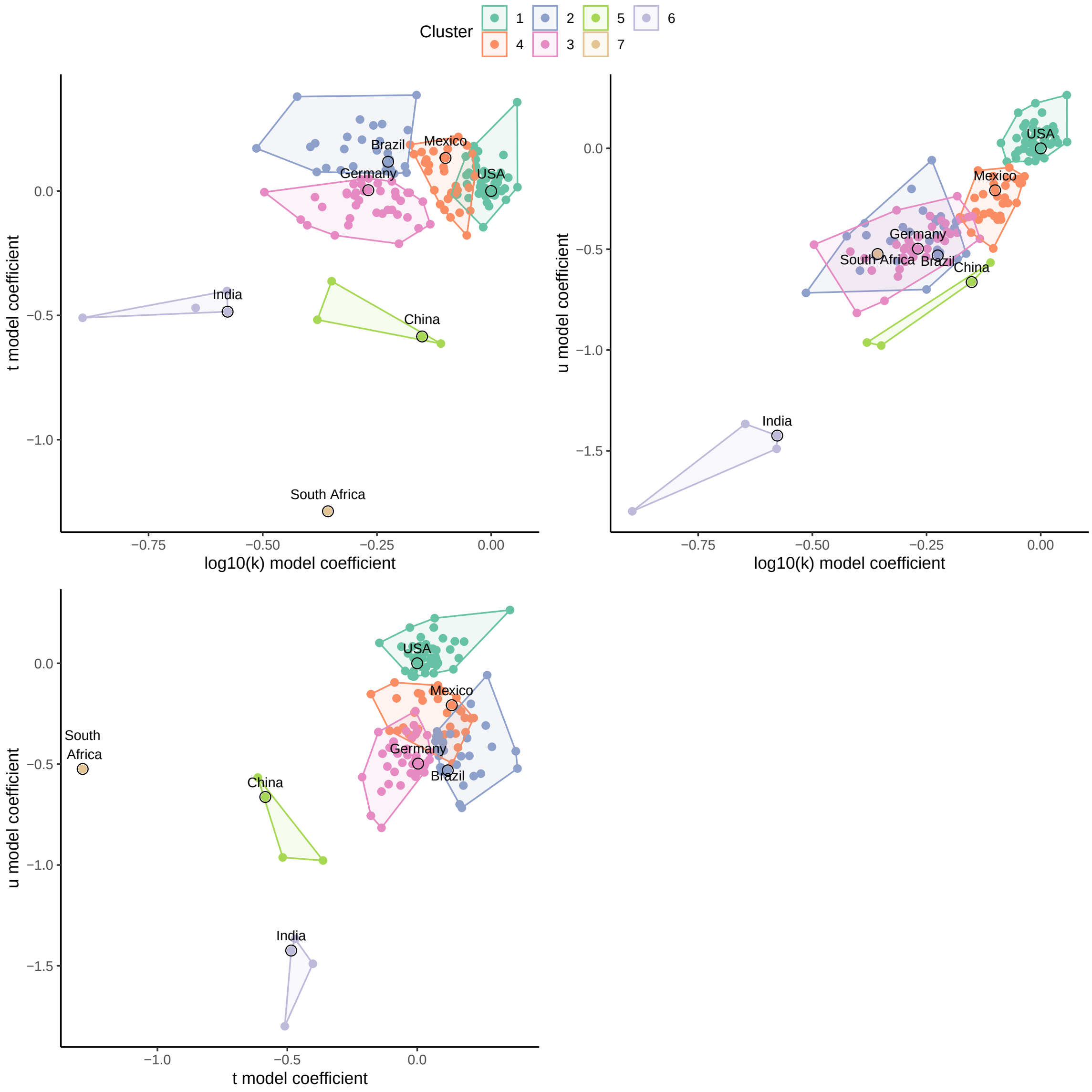

### C Visualization and comparisons of variant transitions and relationships with location characteristics

Figure C.1: Global estimates of $k$, the speed of variant transition. The semi-transparent circles overlaid on the map provide estimates for included sub-national region locations. Some sub-national regions outside of contiguous national boundaries (e.g., Greenland, a sub-region of Denmark) are instead filled in with the appropriate color to reflect the regional value. Countries shown in grey are those for which data were either unavailable or insufficient.

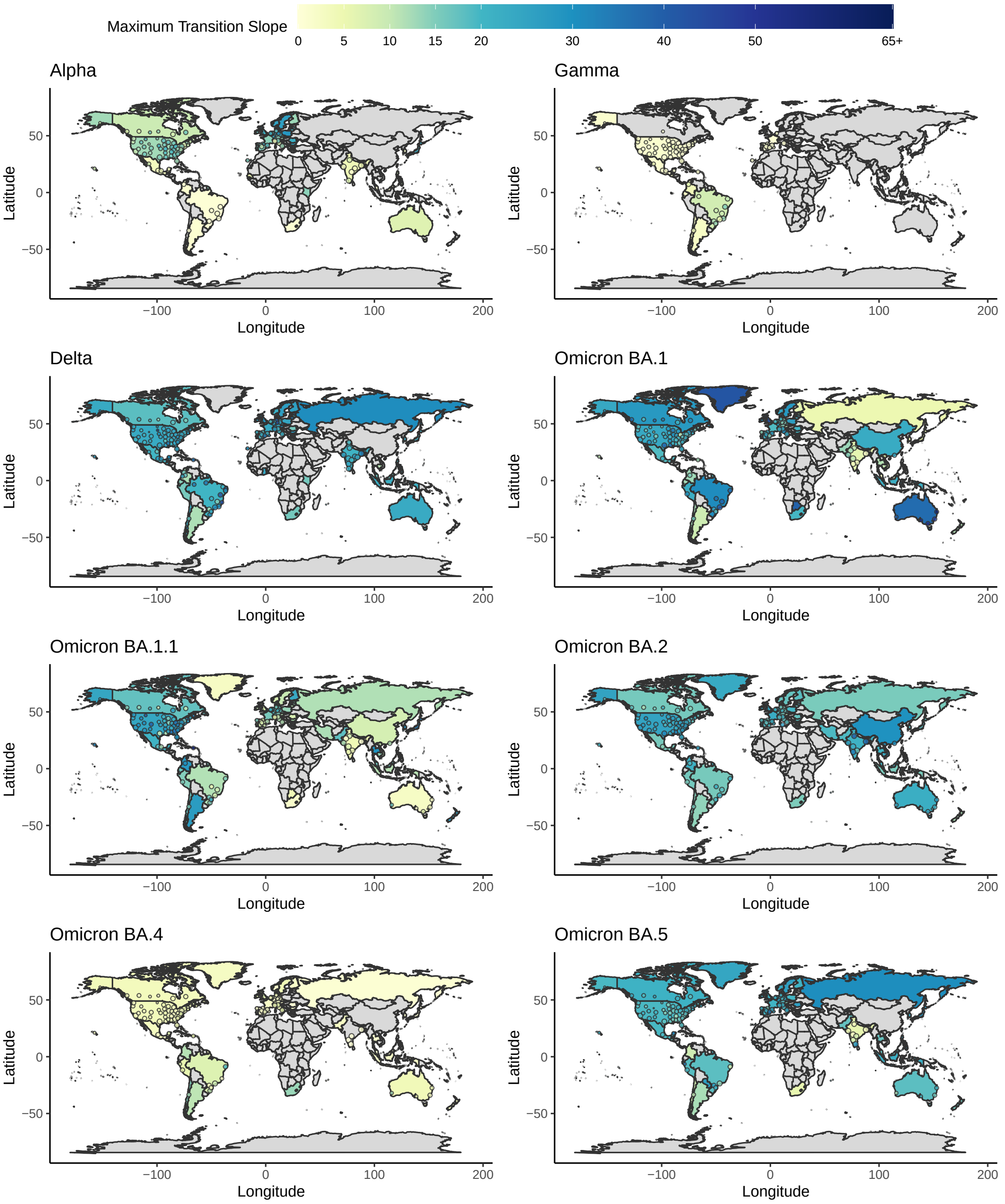

Figure C.2: Global estimates of relative $t0$, the timing of variant transition in days, relative to the first observed value for the variant. See **Supp. Figure C.1** for details.

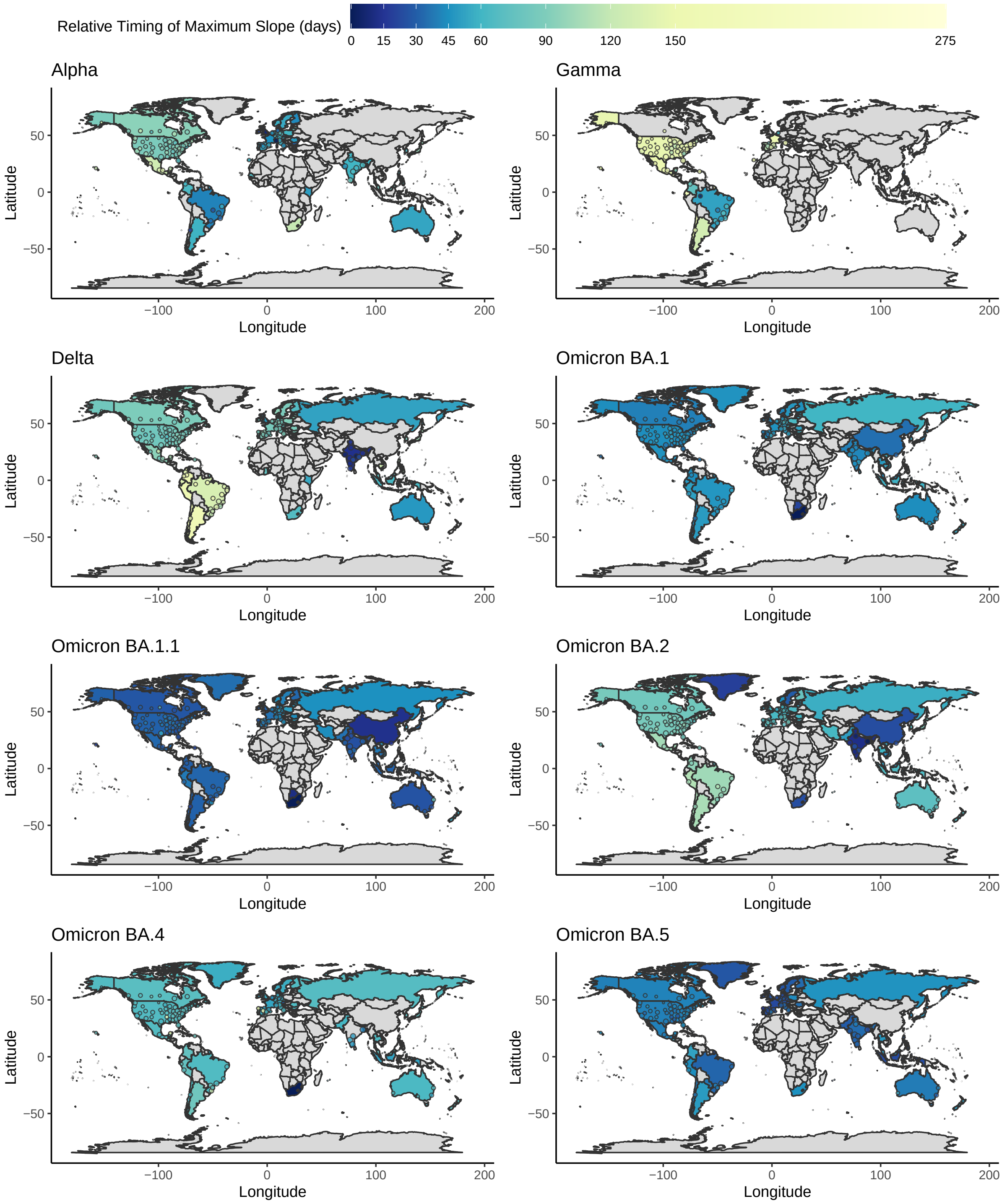

Figure C.3: Global estimates of $u$, the maximum variant prevalence. See **Supp. Figure C.1** for details.

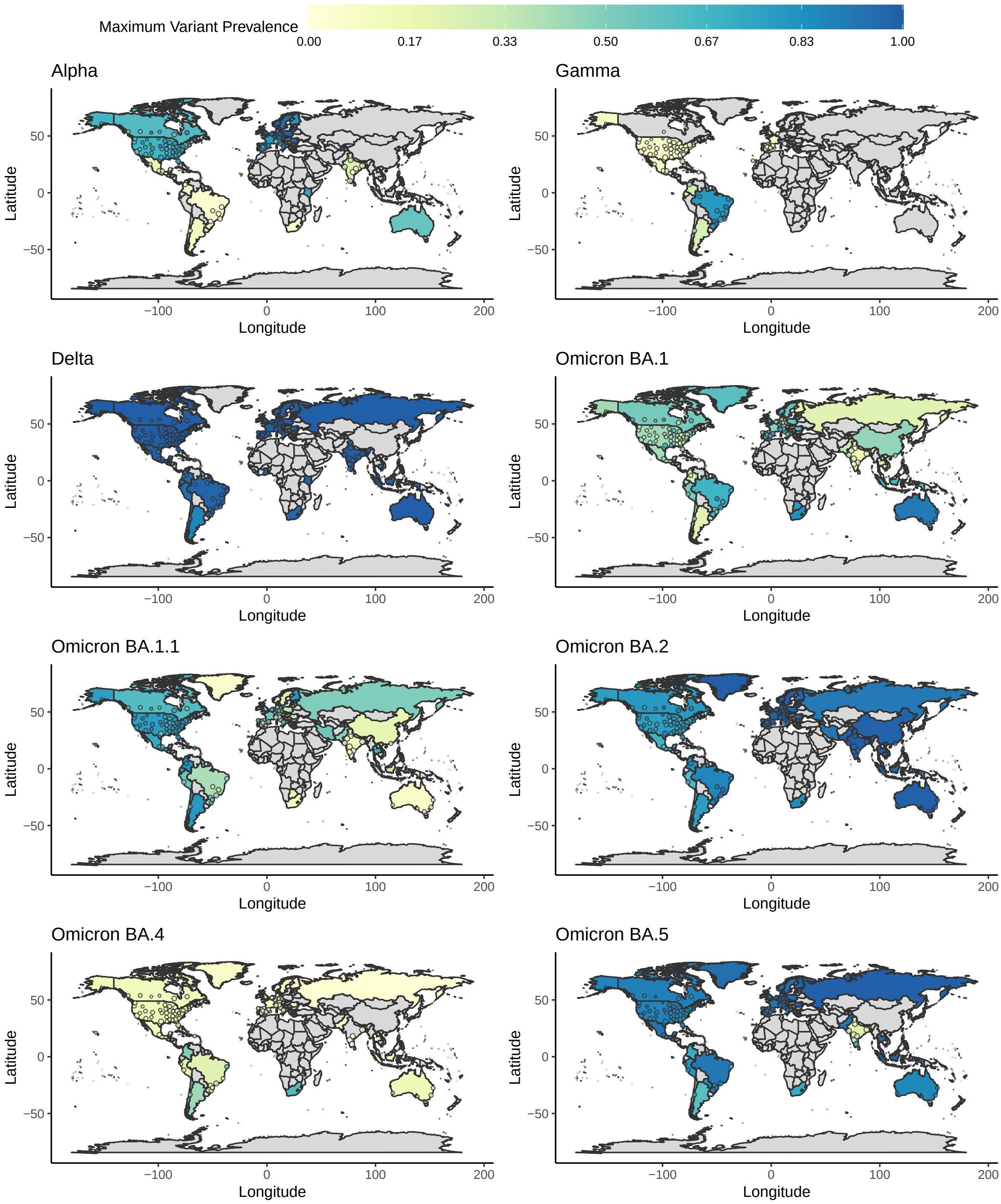

Figure C.4: Cumulative vaccination rates (fully) per 100 over time. The semi-transparent circles overlaid on the map provide estimates for included sub-national region locations. Some sub-national regions are instead shown using filled-in spatial polygons; for example, Greenland is a sub-region of Denmark but shown via polygon in the maps. Countries shown in grey are those for which data were either unavailable or insufficient.

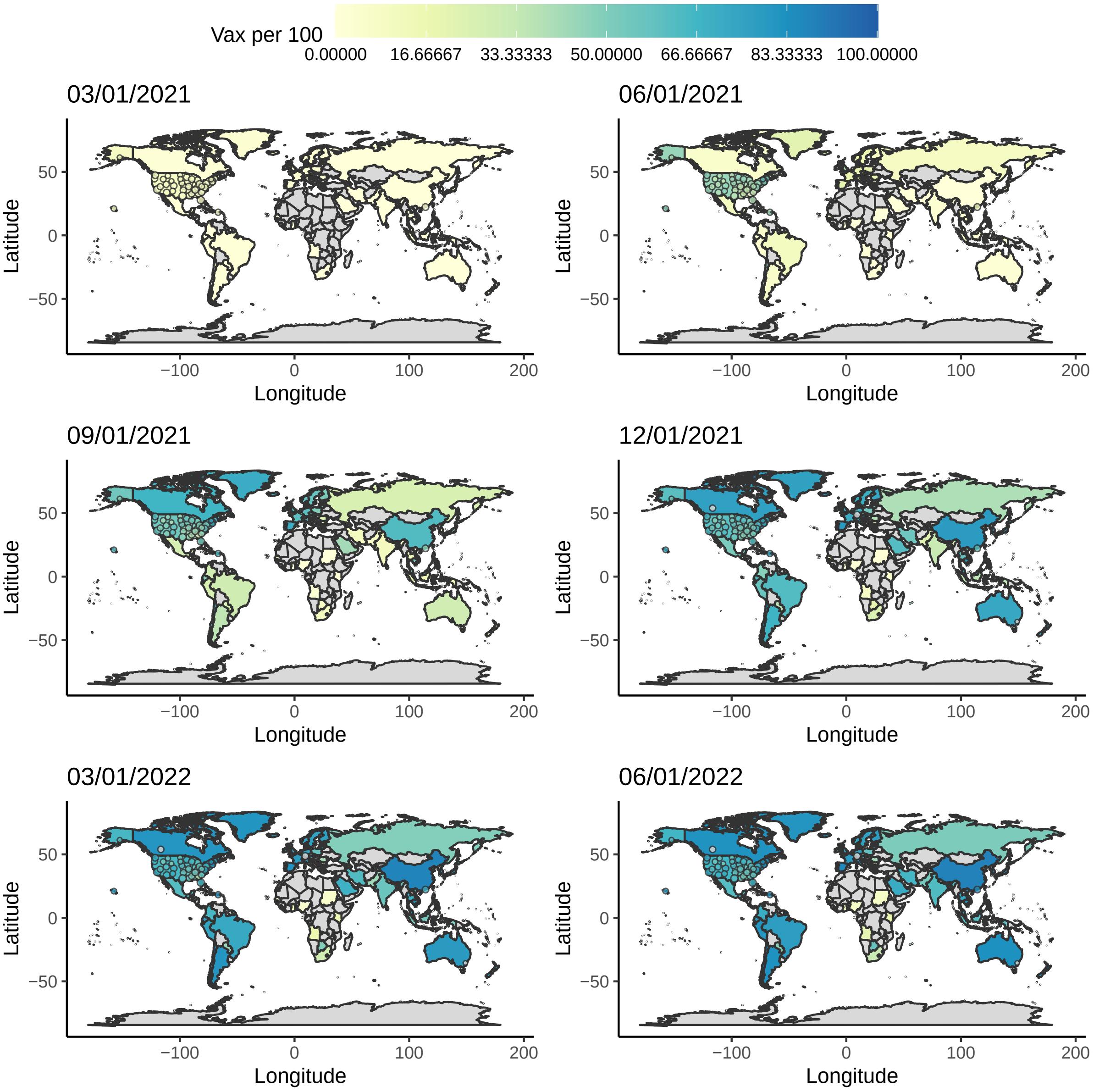

Figure C.5: Spearman correlations between variant transition profile summaries across variants in primary analysis. Numbers in cells reflect the estimated Spearman correlations (among those greater than 0.5)

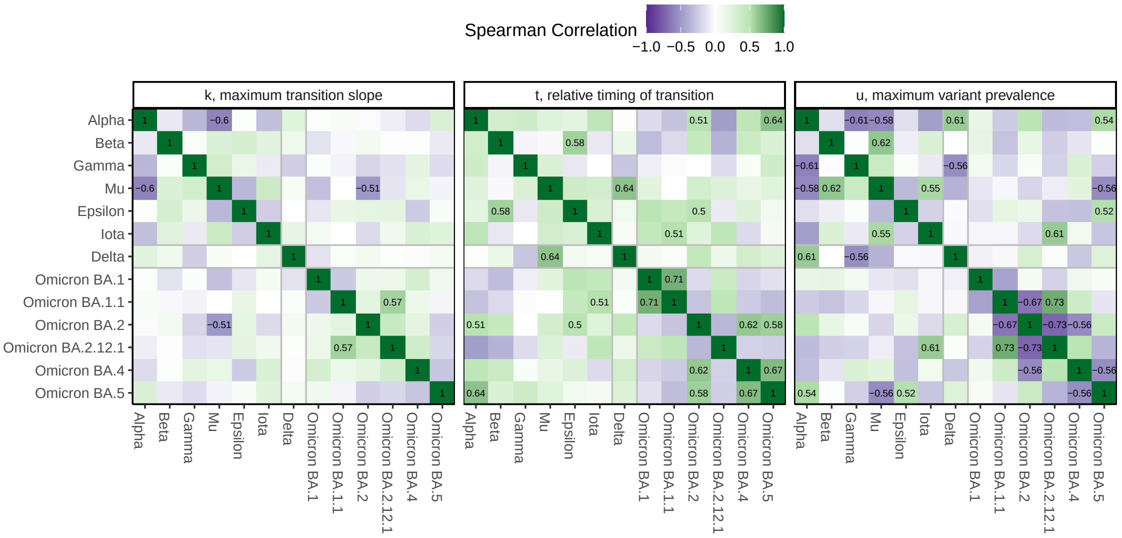


Figure C.6: Spearman correlations between observed location characteristics and variant transition profile summaries by variant in primary analysis. Numbers in cells reflect the number of locations with complete data used for each correlation calculation.

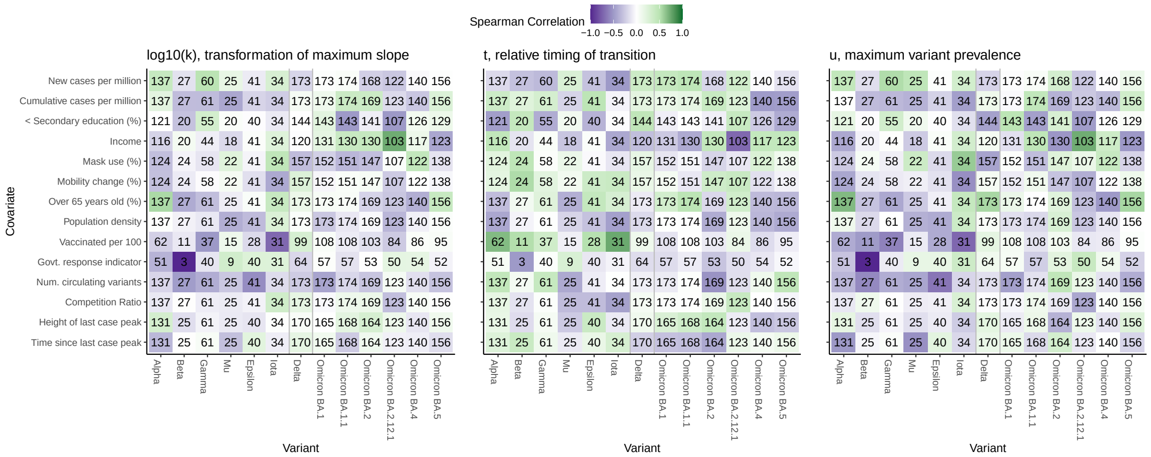


Figure C.7: Kendall's $\tau_b$ correlation between the vaccination rate at the time of variant 5% prevalence andthe timing and magnitude of the maximum slope of the transition curve. Corresponding 95% confidence intervals are also provided. Bolded correlation estimates are significantly non-zero, after adjusting for multiple testing using a Bonferroni correction (30 tests total, including tests for all variants before Mu/Delta and all Omicron sub-variants).

| Variant | Relative Time | Max. Slope |
|---|---|---|
| Alpha | **0.54, ( 0.38, 0.71)** | −0.14, (−0.35, 0.06) |
| Beta | 0.44, (−0.05, 0.93) | −0.15, (−0.60, 0.31) |
| Gamma | 0.27, ( 0.05, 0.50) | **−0.36, (−0.56, −0.16)** |
| Epsilon | **0.37, ( 0.15, 0.58)** | −0.11, (−0.36, 0.13) |
| Iota | **0.55, ( 0.35, 0.75)** | **−0.51, (−0.67, −0.35)** |
| Mu | −0.05, (−0.46, 0.36) | 0.07, (−0.28, 0.41) |
| Delta | 0.08, (−0.07, 0.23) | **0.19, ( 0.08, 0.31)** |
| Omicron BA.1 | −0.10, (−0.25, 0.05) | 0.03, (−0.11, 0.18) |
| Omicron BA.1.1 | 0.01, (−0.14, 0.16) | −0.07, (−0.23, 0.08) |
| Omicron BA.2 | −0.07, (−0.22, 0.07) | −0.10, (−0.24, 0.04) |
| Omicron BA.2.12.1 | 0.10, (−0.05, 0.26) | −0.17, (−0.32, −0.02) |
| Omicron BA.4 | −0.12, (−0.29, 0.04) | 0.04, (−0.13, 0.21) |
| Omicron BA.5 | −0.05, (−0.20, 0.10) | 0.04, (−0.12, 0.20) |

### D Random forest and GLM fit comparisons

**Figure D.1** plots the predicted and multinomial model-estimated ("observed") variant transition summary metrics based on the random forest and GLM regression models. Spearman correlations and Lin's concordance metric are also reported. We see slightly better correlation for the random forest models, which can account for complicated non-linear interactions between location characteristics.

Figure D.1: Comparison between predicted and estimated variant transition summaries from random forest and regression modeling and corresponding Spearman correlations and Lin's Concordance. Higher values of each indicate greater agreement. Spearman correlations are calculated in the *training data* for both models.

(a) Random forest predictions vs. "observed" transition summaries

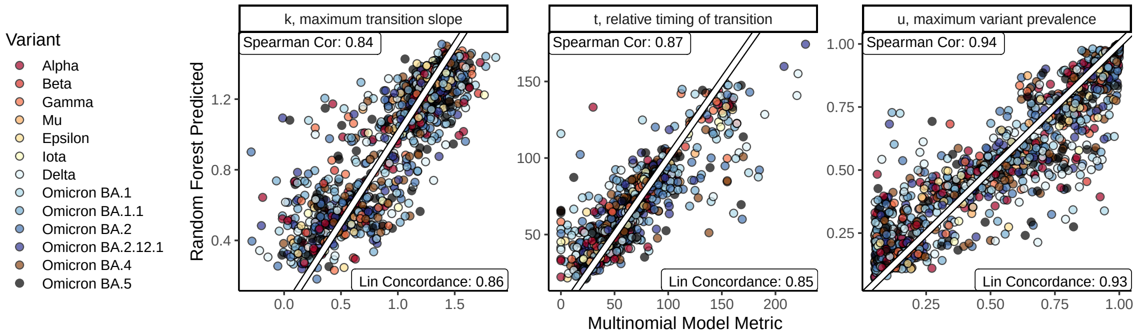


(b) Regression model predictions vs. "observed" transition summaries

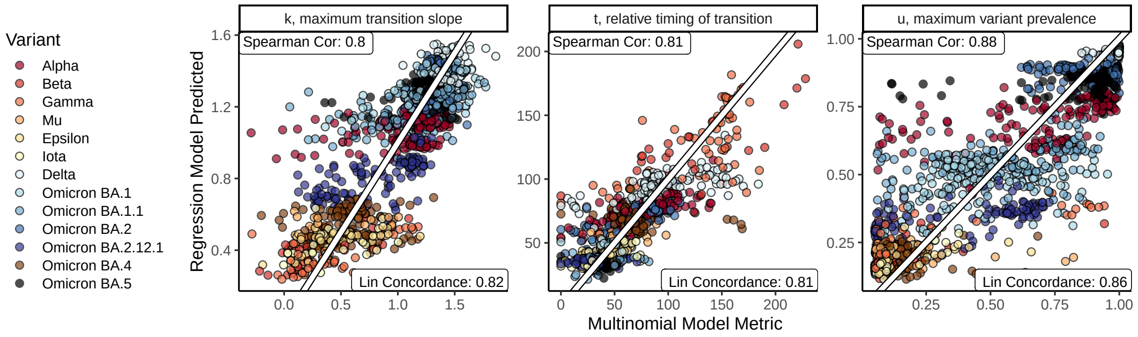


(c) Regression model predictions vs. Random forest predictions

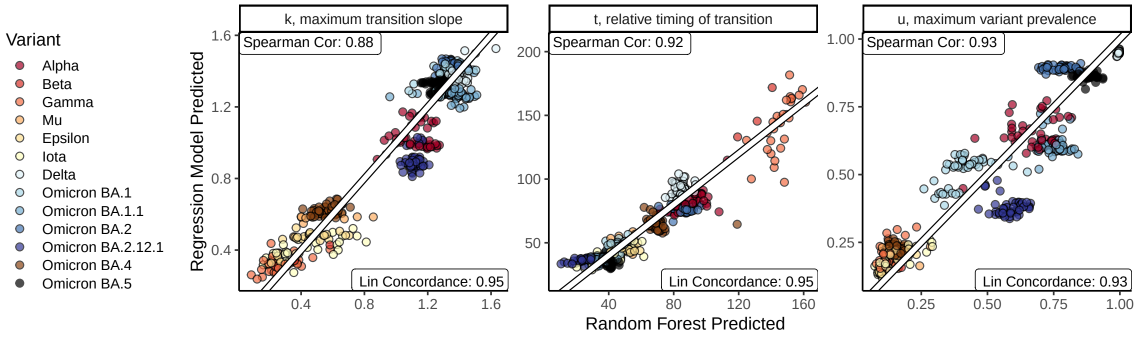

Figure D.2: Relative importance (a) and regression model coefficient estimates (b) of two adjusted models for associations between location characteristics and variant transition summaries, *using only the 479 location and variant combinations with complete data.*[1]

(a) Random forest modeling importance metrics

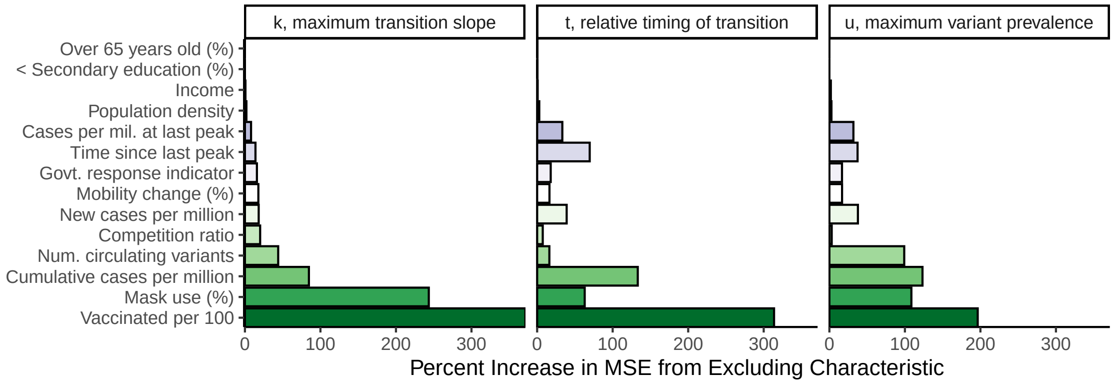


(b) Regression modeling coefficient estimates

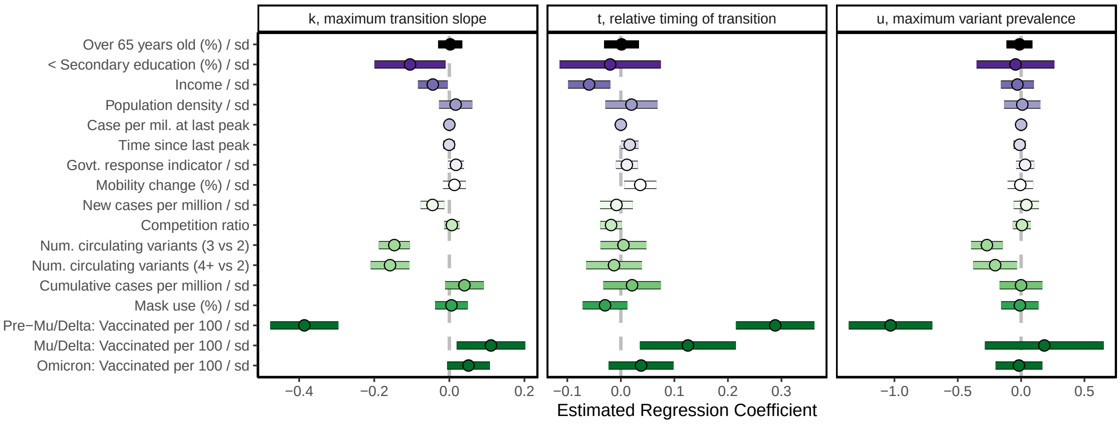


[1] Random forest importance measured in terms of percent increase in mean squared prediction error. For regression modeling, continuous predictors were scaled by their standard deviations. Gaussian, Poisson, and Beta regression were used for $\log_{10}(k)$, $t$, and $u$, respectively. All models also adjusted for variant/sub-variant. Missing predictor information was handled using complete case analysis (i.e., excluding any location x variant combinations with missing predictor information).

[3] WorldPop. URL `www.worldpop.org`.

[4] Shruti Khare, Céline Gurry, Lucas Freitas, Mark B. Schultz, Gunter Bach, Amadou Diallo, Nancy Akite, Joses Ho, Raphael T.C. Lee, Winston Yeo, GISAID Core Curation Team, and Sebastian Maurer-Stroh. GISAID's role in pandemic response. *China CDC Weekly*, 3, 2021. ISSN 20967071. doi: 10.46234/ccdcw2021.255.

[5] IHME. Data and Forecast Repository, 2021. URL `http://www.healthdata.org/covid/data-downloads`.

[6] OECD. Composite leading indicator (CLI). 2021. doi: 10.1787/4a174487-en. (Accessed on 31 August 2021).
